# Structural topology shapes regional roles in neuronal avalanche initiation and termination

**DOI:** 10.64898/2026.10.15.757554

**Authors:** Mario De Luca, Matteo Demuru, Emahnuel Troisi Lopez, Enrica Gallo, Gianmarco Cafaro, Gian Marco Duma, Damien Depannemaecker, Gabriele Casagrande, Mario Quarantelli, Carmine Granata, Giuseppe Sorrentino, Pierpaolo Sorrentino, Marianna Angiolelli

**Affiliations:** Department of Medical, Motor and Wellness Sciences, University of Naples “Parthenope”, 80133 Napoli, Italy; Department of Education and Sport Sciences, Pegaso Telematic University, 80143, Naples, Italy; Department of Physics “E.R. Caianiello”, University of Salerno, Fisciano, Italy; Epilepsy and Clinical Neurophysiology Unit, IRCCS Eugenio Medea, Conegliano, Italy; Aix-Marseille Univ, INSERM, INS, Institut de Neurosciences des Systèmes, Marseille, France; Institute of Biostructures and Bioimaging, Italian National Research Council, Naples, Italy; Institute of Applied Sciences and Intelligent Systems of National Research Council, Pozzuoli, Italy

**Keywords:** Magnetoencephalography data, Neuronal avalanches, Brain network, Graph theory

## Abstract

Whole-brain dynamics unfold as fast, discrete, transient events known as neuronal avalanches, whose propagation is constrained by the connectome, that is, the large-scale scaffolding of structural connections wiring the brain. However, the role of the connectome in determining the topographies of the start and end of these transient cascades remains unclear. Leveraging resting-state Magnetoencephalography (MEG) recordings and diffusion magnetic resonance imaging (dMRI) tractography across 42 healthy individuals, we examined whether the structural architecture of white matter tracts dictates the spatial origins and the end of cortical avalanches. We found that the probability of triggering and finishing an avalanche is strongly related to the structural nodal strength, showing a positive correlation that is highly robust to non-parametric spatial permutation testing and bootstrapping. This structure-function coupling is highly stable and robust across different activation thresholds, demonstrating that the choice of avalanche extraction threshold does not drive or alter our main findings. Collectively, our results suggest that structural properties determine a region’s propensity to initiate and terminate neuronal avalanches.

## 1. Introduction

Neuronal avalanches describe spontaneous brain activity in which transient cascades propagate through distributed networks ^1^. Converging evidence indicates that these cascades are constrained by the underlying anatomical scaffold: in humans, the structural connectome strongly shapes the fast spreading of neuronal avalanches, spatially ^2^ and temporally ^3^. Furthermore, computational modelling suggests that structural heterogeneity and network topology can regulate where and when avalanches are triggered ^4^. Neuronal avalanches can be understood as an emergent expression of the brain’s multiscale organization. Fast microscopic events are organized, generating collective wave-like activities that spread temporally and spatially at the scale of the whole brain. In particular, sizes and durations of these non-linear bursts follow characteristic scale-free laws ^5,6^. At the whole-brain level, this recruitment is constrained by the local dynamical state of individual regions and the hierarchical architecture of the structural connectome, which imposes a repertoire of intrinsic modes ^7^. The large-scale structural connectome, however, is far from homogeneous, as it displays both scale-free and small-world properties ^8^. This topological heterogeneity across regions suggests that the ability to ignite neuronal avalanches might not be uniformly distributed but constitutes a region-specific property. Whole-brain modelling provides direct mechanistic support for this hypothesis. Castro et al. showed that the empirical human connectome enables a small, compact set of regions to enter a self-sustained high-activity state at substantially lower levels of global excitation than randomized connectomes ^9^. These first-ignited regions corresponded to the strongest weighted core of the connectome, while more regions are subsequently recruited as a function of global excitability. This large-scale organization echoes observations at the cellular level, where population bursts are consistently preceded by a restricted set of relatively stable “leader” neurons. The early recruitment of these neurons predicts whether a local precursor will develop into a full burst, while leader identity carries information about the ensuing pattern of network activation, suggesting that they act as nucleation sites ^10^. Consistently, simulations of sparse neuronal networks show that highly connected neurons tend to activate earlier during an avalanche ^11^. Taken together, these findings suggest that the capacity to initiate and support large-scale activity is determined by a node’s embedding within a wider network architecture. By extension, the specific topological position of each cortical region may confer a distinct propensity to initiate and sustain the recruitment of the surrounding network, acting as an “avalanche starter” or “avalanche ender”.

In this study, we hypothesized that a cortical region’s position within the structural connectome and its local topological organization shape its propensity to initiate and support large-scale activity cascades. Specifically, we tested whether regions with stronger structural embedding, indexed by nodal strength, or greater topological centrality were more likely to initiate and sustain neuronal avalanches throughout. To this end, we combined source-reconstructed magnetoencephalography recordings with diffusion MRI-derived (dMRI) tractography from healthy participants, previously published in (CIT 2). We extracted neuronal avalanches from source-reconstructed MEG data acquired in 42 healthy participants and, for each region, quantified its conditional probabilities of acting as an avalanche starter or ender, defined as the proportions of avalanches involving that region in which it was active in the first or final time bin, respectively. We correlated the region’s propensities to act as a starter or ender with their corresponding centrality in the structural network at both the whole-group and individual levels.

## 2. Materials and Methods

### 2.1 Participants

The study cohort consisted of 42 healthy adult participants (25 males, 17 females; mean age 26.7 ± 5.6 years). Eligibility required that individuals be free from significant internal, psychiatric, or neurological conditions and abstain from any medications known to influence MEG or MRI data quality. This research adhered to the ethical standards outlined in the Declaration of Helsinki, with oversight and approval provided by the “Comitato Etico Campania Centro’’ (Prot.n.93C.E./Reg. n.14-17OSS). Each participant provided formal written consent before enrollment.

### 2.2 MEG preprocessing and source reconstruction

Our recordings were captured using a 163-magnetometer system engineered by the National Research Council of Italy, Institute of Applied Sciences and Intelligent Systems (ISASI). The acquisition protocol consisted of two consecutive 3.5 minute sessions while the participants maintained a resting state with eyes closed. Head positioning was tracked by digitizing four anatomical landmarks alongside four auxiliary coils, while cardiac and ocular activity were simultaneously monitored via ECG and EOG. Raw signals were sampled at 1024 Hz following anti-aliasing filtration, after which a fourth-order Butterworth IIR band-pass filter (0.5–48 Hz) was utilized. Environmental interference was mitigated through principal component analysis of data from reference sensors. Subsequently, supervised independent component analysis was applied to isolate and suppress physiological noise. Finally, a specialist visually inspected the data to exclude compromised channels. We performed source reconstruction across 116 regions of interest (ROIs) defined by the AAL atlas ^12,13^. We implemented the volume conduction model and utilized the linearly constrained minimum variance (LCMV) beamformer ^14^, integrating native structural MRI data for each participant. Source activity was estimated specifically at the centroid of each defined region. Following initial processing, we restricted our analysis to 78 AAL regions, omitting the 26 cerebellar ROIs due to inherent limitations in MEG reliability within those areas and the 10 ROIs of subcortical areas, keeping only the cortical regions. All data transformation and source estimation procedures were executed using the Fieldtrip analysis environment ^15^. Resting-state MEG recordings were structured in Brain Imaging Data Structure (BIDS) format. The epochs of the signal classified as “good” through visual inspection were converted to arrays via MNE-Python, concatenated, and then z-scored point-by-point along the time axis. To prevent statistical bias from varying recording lengths, an algorithm identified the global minimum duration across the cohort and uniformly truncated all subject data matrices to this boundary length before analysis.

### 2.3 MRI acquisition

High-resolution anatomical and diffusion MRI (dMRI) data were acquired for each participant using a 1.5 Tesla scanner (Signa, GE Healthcare) following the MEG acquisition session. T1-weighted 3D structural volumes were gathered using a Magnetization-Prepared Gradient-Echo (BRAVO) sequence with the following parameters: TR/TE/TI = 8.2/3.1/450 ms, isotropic voxel size = 1 × 1 ×1 mm^3^, 50% partition overlap, and 324 sagittal slices providing full brain coverage. Diffusion-weighted images were collected using an Echo-Planar Imaging (EPI) sequence (TR/TE = 12,000/95.5 ms, voxel resolution = 0.94 × 0.94 × 2.5 mm^3^) along 32 diffusion-sensitizing directions, complemented by 5 unweighted (B_0_) volumes. Preprocessing of the dMRI datasets was performed using tools from the FMRIB Software Library (FSL). Motion artifacts and eddy current-induced distortions were corrected via the eddy_correct tool, with appropriate reorientation of the diffusion gradient vectors. Brain tissue was isolated from non-brain structures using the Brain Extraction Tool (BET) applied to B_0_ acquisitions. A diffusion tensor model was fitted at each voxel to estimate local diffusion orientation. Whole-brain streamline reconstruction was executed using deterministic tractography (Diffusion Toolkit) based on the Fiber Assignment by Continuous Tracking (FACT) algorithm, implementing a maximum turning angle threshold of 45°, spline filtering, and a fractional anisotropy (FA) termination threshold of 0.2. To map structural connectivity onto the functional parcellation, the gray matter regions of interest (ROIs) from the Automated Anatomical Labeling (AAL) atlas were transformed into each subject’s native space. Specifically, individual FA maps were normalized to the MNI-space FA template via SPM12 spatial normalization routines. The inverse transformation matrices were then applied to the AAL ROIs—masked by a gray matter probability threshold of 0.2—mapping them onto native structural spaces, with accurate alignment verified through visual inspection. Finally, structural connectivity matrices were built by calculating the total number of interconnecting streamlines between each pair of cortical AAL ROIs using custom scripts in Interactive Data Language (IDL). Brain space was parcellated into 78 regions following the Automated Anatomical Labeling (AAL) atlas. A symmetric 78×78 structural matrix was generated for each subject, and a group-level average matrix was derived.

### 2.4 Avalanche extraction and “avalanche starter” and “avalanche ender” identification

Neuronal avalanches were identified from source-reconstructed brain activity after z-score normalization of each regional time series. The detection threshold was selected using a data-driven procedure based on the statistical properties of the signals. For each subject, the normalized time series from all brain regions were concatenated to obtain a single empirical amplitude distribution, which was then compared with the Gaussian distribution expected for background fluctuations. The threshold was chosen as the amplitude at which the empirical distribution began to systematically deviate from Gaussian behavior, indicating the emergence of rare, large-amplitude events that are unlikely to arise from stochastic fluctuations alone ^16^. The rationale behind this approach is that low-amplitude fluctuations are approximately Gaussian, whereas the heavy tails of the empirical distribution reflect coordinated neuronal activity. The main analyses were performed using an activation threshold of |z| > 2.5. To assess sensitivity to this choice, the complete avalanche extraction and analysis pipeline was repeated at |z| > 2.0 and |z| > 3.0. An avalanche was defined as a continuous period during which at least one brain region exhibited activity above the selected threshold (Fig. 1a). More specifically, an avalanche started when the first suprathreshold event appeared in any region and ended when the activity of all regions simultaneously returned below threshold, ensuring that each avalanche was preceded and followed by a silent interval (Fig. 1b). Only avalanches lasting at least two time bins were included. As an additional robustness analysis, we repeated the analyses after further restricting the retained avalanches to those recruiting at least two distinct brain regions over their duration. This criterion excludes events confined to a single region, allowing us to assess whether the findings persist when considering only avalanches involving spatial propagation across regions. Regional starter and ender probabilities were recomputed using only the avalanches satisfying this additional criterion.

**Figure 1.**
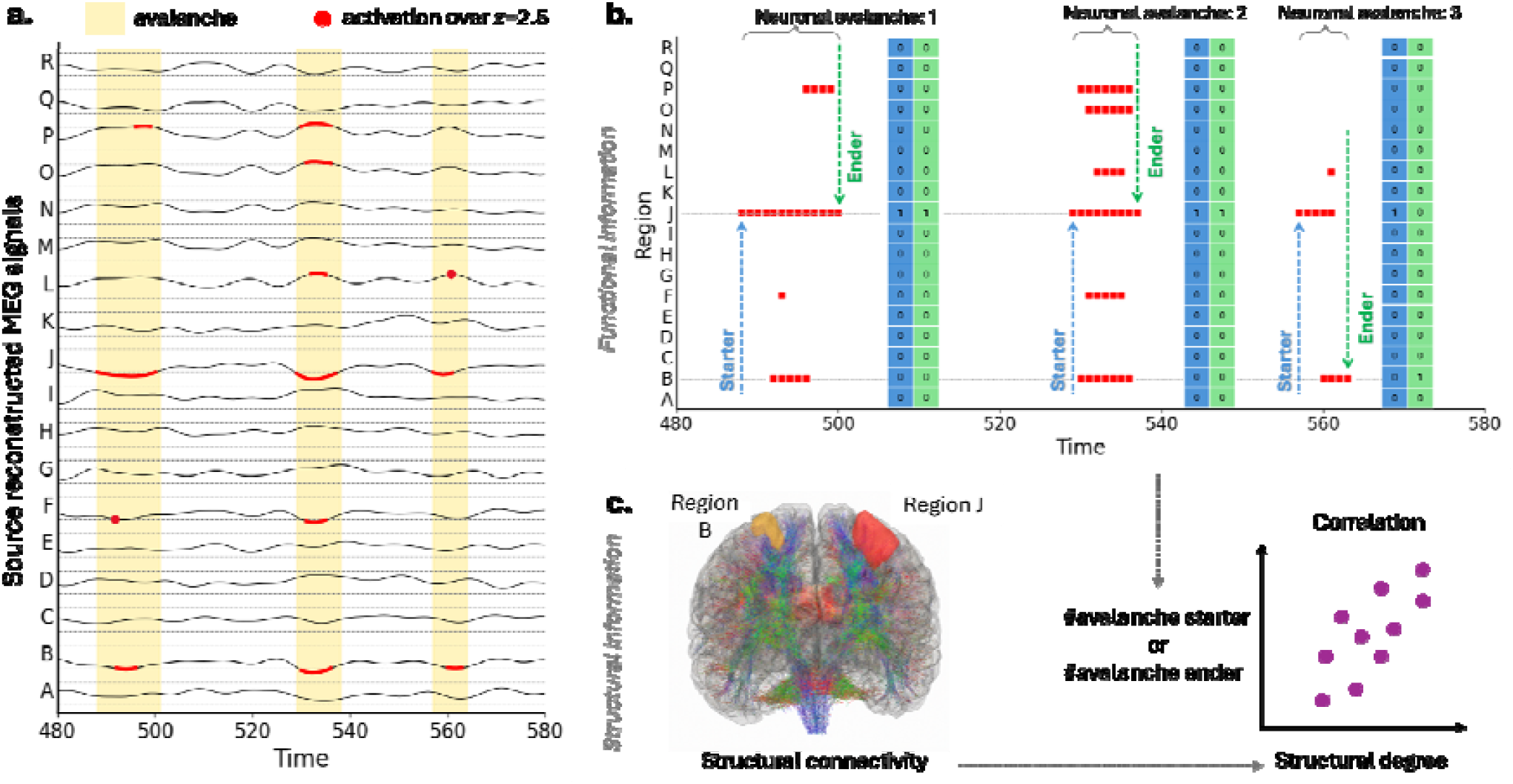
**(a)** Identification of neuronal avalanches and extraction of regional dynamical features to investigate how the propensity of brain regions to initiate and end neuronal avalanches is constrained by the underlying structural connectome. Continuous source-reconstructed (using AAL Atlas) brain activity is first thresholded to identify neuronal avalanches, defined as uninterrupted periods of suprathreshold activity (yellow shaded windows). Within each avalanche, the temporal evolution of regional activations is represented in a binary raster (top right), where each red dot denotes a suprathreshold activation of a brain region at a given time bin. **(b)** For every avalanche, regions active in the first time bin are labelled as avalanche starters (blue), whereas regions active in the final time bin are labelled as avalanche enders (green). These labels are encoded as binary vectors and accumulated across avalanches to quantify, for each region, its propensity to initiate or terminate large-scale activity cascades; that is, the fraction of avalanches in which the region is classified as a starter or an ender. **(c)** Finally, the extracted functional descriptors are integrated with the structural connectome to investigate whether the regional tendency to initiate or terminate neuronal avalanches is constrained by the underlying anatomical network topology. The resulting framework enables the systematic characterization of the relationship between local structural connectivity and the dynamical role played by each brain region during large-scale spontaneous activity propagation.

To characterize the temporal organization of avalanche propagation, we identified the regions responsible for initiating and terminating each avalanche. For each brain region, its tendency to initiate avalanches is quantified by computing the *starter frequency*, defined as:

where is the number of avalanches in which i-region acted as a starter and is the total number of avalanches in which the same region was recruited at any point during the avalanche ^17,18^. Similarly, the *ender frequency* was defined as:

where denotes the number of avalanches in which region (i) was active during the final time bin.

The statistical characterization of avalanche sizes in a subset of the same source dataset has been reported in (Cafaro et al., 2026) ^19^, where distinct size regimes were identified and truncated power-law scaling was reported in the large-avalanche regime, with an exponent close to the mean-field prediction of 1.5. This previous characterization supports interpreting the extracted sequences as neuronal avalanches, while the present study focuses on their regional initiation and termination patterns.

### 2.5 Structural connectome metrics

To characterize the topological properties of the structural connectome, the Brain Connectivity Toolbox for undirected weighted networks has been used ^20^. The metrics used in this study are the nodal strength, the betweenness centrality, and the eigenvector centrality.

In a structural brain network, where nodes represent gray matter regions and edges represent white matter tracts, the nodal strength is the sum of the weights of all connections attached to a single node. Structural nodal strength was formally defined as:

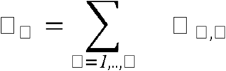

Where □ _□, □_ is the weight of the connection between node *i* and node *j*.

Betweenness centrality measures how frequently a node lies on the shortest paths connecting pairs of other nodes in the network. Formally, the betweenness centrality g(v) of a node v is defined as:

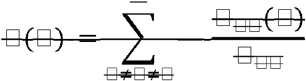

where □ _□ □_ □ is the total number of shortest paths from node □ to node □ and □ _□ □_ (□) is the number of those paths that pass through □ (not where □ is an endpoint).

Let □ = (□, □) be a graph with adjacency matrix □ = (□ _□, □_), where □ _□, □_ = *1* if node □ is linked to node □, and □ _□, □_ = *0* otherwise. The eigenvector centrality score □ _□_ of a node □ is defined as:

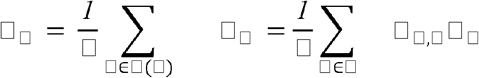

Where □ (□) is the set of neighbors of node □; □ is a constant (specifically, the largest eigenvalue); and □ _□_is the centrality score of neighbor node t.

A node scores highly if it is linked to other highly connected, central regions, reflecting its importance in the global network architecture.

### 2.6 Statistical Analysis

We quantified the coupling between structural measures and functional initiation and finishing probability using Spearman correlation. We performed non-parametric permutation tests by shuffling the order 10,000 times to generate a surrogate null distribution and assess observed significance. Specifically, by randomly shuffling the spatial region labels of the structural strength vector, we disrupted the connectome’s empirical topology while preserving the statistical distribution of the degrees.

To assess the stability and robustness of the structure-function correlation, we employed a bootstrapping approach. We generated 5000 bootstrap samples by resampling with replacement from the original cohort of 42 participants. For each bootstrap sample, we recomputed the Spearman correlation tests between structural nodal strength and avalanche starter and ender frequency. This allowed us to estimate the 95% confidence interval (CI) for the correlation coefficient, ensuring the observed relationships were not driven by outliers or specific subjects.

## 3. Results

### 3.1 The structural connectome shapes both starter and ender propensities of neuronal avalanches

We examined the relationship between regional structural strength and the propensity of brain regions to participate in neuronal avalanche onset and termination in 42 healthy participants. Avalanches were identified in source-reconstructed MEG time series across 78 regions using a threshold of |z| > 2.5, retaining events lasting at least two time bins. For each region, starter and ender frequencies were defined as the proportion of avalanches recruiting that region in which it was active in the first or last time bin, respectively.

At the group level, regional structural strength was positively correlated with mean starter frequency (Spearman’s ρ = 0.340, p = 0.0023; Fig. 2a) and mean ender frequency (ρ = 0.353, p = 0.0015; Fig. 2b). Thus, regions with greater structural strength were more likely to be active at both avalanche onset and termination. The linear regression lines illustrate the positive trends in the regional measures, whereas the reported coefficients quantify their rank associations. The null-distribution panels (Fig. 2c, 2d) compare the observed structural–functional correlations with their corresponding null distributions. For both starter and ender frequencies, the observed correlations, indicated by red horizontal lines, lie in the upper tails of distributions concentrated near zero. These comparisons provide additional evidence that the positive regional associations are unlikely under the null models used. The bootstrap panels display the resampling distributions and 95% CI of the starter–strength [0.289 - 0.398] and ender–strength [0.283 - 0.402] correlations. Both distributions are concentrated around positive values, with the observed coefficients close to their centers. The shaded intervals also lie above zero, supporting the stability of the positive associations under the bootstrap procedure. On the other hand, betweenness centrality showed (not reported here) no comparable association with starter and ender frequencies (p = 0.47 | p = 0.06).

**Figure 2.**
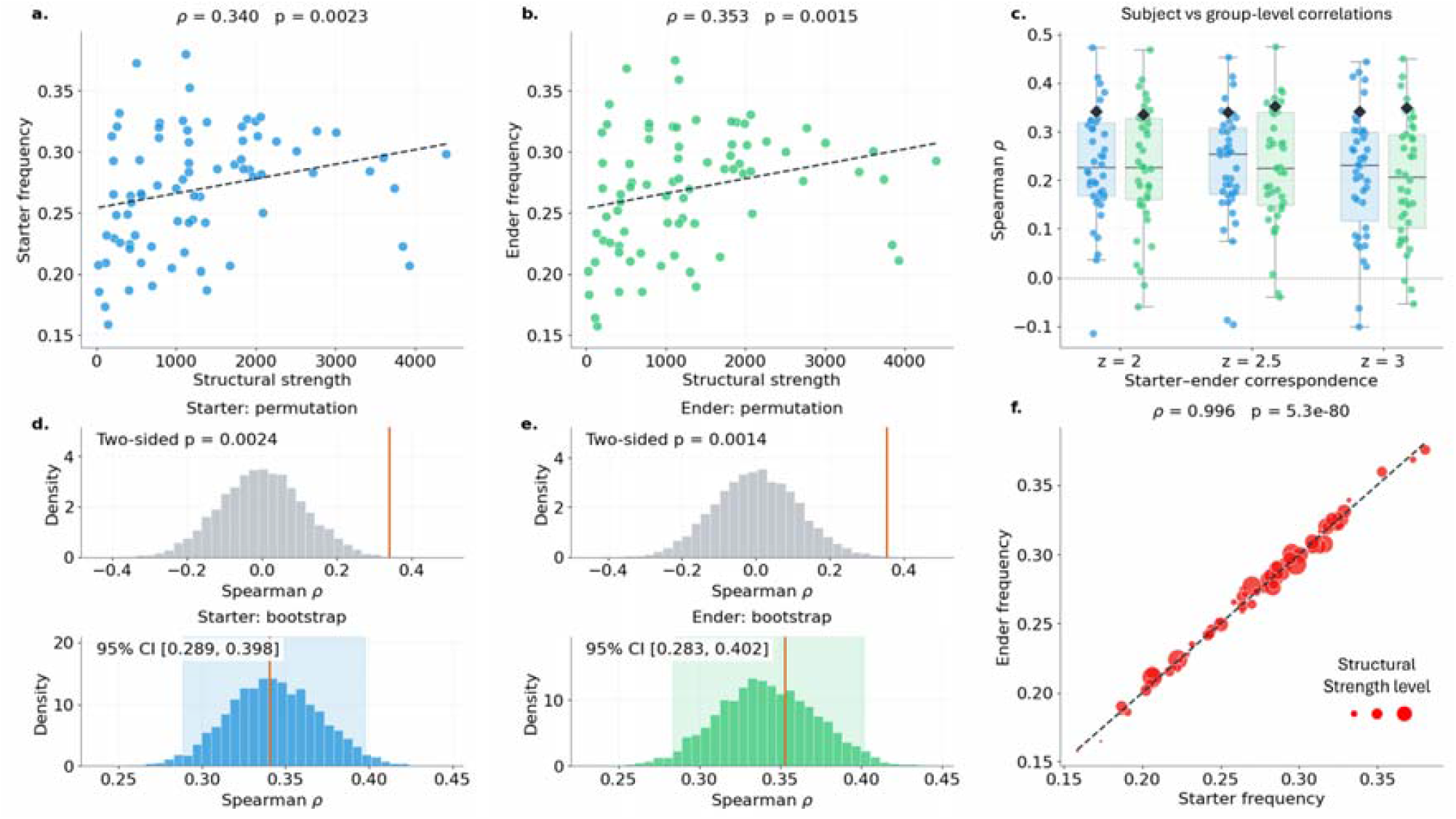
Structural connectivity and regional participation in neuronal avalanche onset and termination. (**a, b**) Group-averaged conditional starter and ender frequencies as a function of regional structural strength. Lines represent ordinary least-squares fits; annotations report Spearman’s rank correlations and their associated p-values. (**c, d**) Null distributions obtained by permuting regional strength labels and bootstrap distributions obtained by resampling participants at z = 2.5. Vertical red lines indicate observed correlations; shaded bands represent 95% percentile bootstrap confidence intervals. (**e**) Subject- and group-level correlations across three activity thresholds; diamonds indicate correlations between group-averaged regional frequencies and structural strength. (**f**) Relationship between regional starter and ender frequencies, with the size proportional to structural strength. Starter and ender frequencies are conditional on regional recruitment.

Fig. 2e compares subject-level and group-level correlations for starter and ender frequencies at three different thresholds (|z| = 2, 2.5, and 3) used to detect neuronal avalanches. Correlations were positive in most participants for both measures at each threshold, although their magnitude varied across individuals, and a small number of participants exhibited negative correlations. We calculated group-level correlations after averaging regional starter and ender frequencies across participants. The group-level coefficients (black diamonds) exceeded the mean individual-level coefficients across all three thresholds. This pattern suggests that the regional association becomes more pronounced after averaging across participants, potentially reflecting an attenuation of individual measurement noise. Complementing these findings, Fig. 2f directly compares the regional starter and ender frequencies and shows that starter and ender frequencies were strongly positively correlated across regions (ρ = 0.996, p = 5.3 × 10□ □ □). Regions with higher starter frequencies also exhibited higher ender frequencies, revealing closely aligned regional profiles of participation in avalanche onset and termination. The point size represents regional structural strength, allowing this anatomical measure to be visualized alongside the relationship between the two frequencies.

These findings were also supported when the analyses were restricted to avalanches recruiting at least two distinct regions. At the main activation threshold of |z| > 2.5, structural strength remained positively associated with starter frequency (Spearman’s ρ = 0.316, p = 0.0049) and ender frequency (ρ = 0.296, p = 0.0086). Both associations were supported by two-sided permutation tests (p = 0.0056 and p = 0.0079, respectively), and their 95% bootstrap confidence intervals excluded zero (starter: [0.224, 0.376]; ender: [0.207, 0.377]). Positive group-level associations were retained across all three activation thresholds, and regional starter and ender frequencies remained strongly correlated (ρ = 0.980, p = 3.0 × 10□ □ □).

## 4. Discussion

In this manuscript, we characterized the relationship between a region’s topological role within the structural connectome and its ability to initiate and end large-scale neuronal avalanches. Using structural dMRI data and source-reconstructed MEG recordings, we showed that a region’s structural nodal strength is strongly and directly related to how often it initiates and ends neuronal avalanches. This strong positive correlation supports a “hub-centric” model of dynamic activity generation. Regions with high structural strength—that is, a high number of structural connections—may be better positioned to collect and integrate activity from other areas. Along these lines, we propose that activity propagating through the brain is more likely to converge on and reverberate within central regions, facilitating avalanche initiation and helping these regions sustain suprathreshold activity throughout the avalanche.

These findings provide an empirical counterpart to previous whole-brain modeling work showing that the interaction between regional dynamics and long-range structural coupling is predicted to induce regional differences in ignition propensity ^9^. Castro et al. demonstrated in silico that regions belonging to the maximally strong weighted core of the human connectome were the first to sustain an ignited state as interregional coupling increased. Moreover, the empirical human connectome required less global coupling to initiate ignition than randomized surrogate networks and supported a more gradual recruitment of additional regions. Its weighted core-shell organization therefore provided a compact substrate in which strong mutual interactions could both facilitate ignition and stabilize the resulting high-activity state. Although the bistable states examined in that model differ from the short-lived neuronal avalanches studied here, both the previous modeling results and our empirical findings support a common organizational principle: the structural connectome not only constrains the propagation of activity once initiated ^2^ but also shapes regional differences in the likelihood that a local fluctuation will evolve into a large-scale event. This interpretation is consistent with broader evidence that connectome topology shapes cooperative spreading and the recruitment of distributed cortical regions ^21^. Furthermore, the connectome may influence cascades at multiple spatial scales, linking the locations of start and finish of neuronal avalanches to frameworks such as intrinsic ignition and interareal amplification ^22,23^.

The lack of comparable association of the betweenness centrality with avalanche initiation or termination, suggesting that occupying a strategic position along the shortest communication paths is not, by itself, relevant to ignite one region or sustain its activation throughout a burst. By contrast, an association with eigenvector centrality (not reported here) emerged after log-scaling the structural weights. The relationships of avalanche initiation and termination with nodal strength and eigenvector centrality may not reflect independent topological mechanisms. In heterogeneous networks, the components of the principal eigenvector can be approximately proportional to node degree under mean-field conditions, whereas in other regimes eigenvector centrality becomes concentrated around the largest hubs or densely connected cores ^24,25^. Our findings may be reminiscent of this coupling between local connectivity and spectral centrality. By compressing the influence of the strongest connections, logarithmic rescaling may provide an eigenvector-centrality estimate that is less dominated by a small number of extreme weights and more sensitive to a region’s broader embedding. From this perspective, the association with avalanche initiation and termination could indicate that cascade onset is favored not only by large structural connections, as captured by nodal strength, but also by neighboring structurally influential regions. However, this interpretation remains tentative.

Furthermore, the remarkable stability of this structure-function coupling across different signal binarization thresholds highlights the physiological and methodological robustness of our findings, which are unlikely to be a statistical artifact of a specific signal binarization. While beyond the scope of this manuscript, avalanches have been shown to preserve statistical properties across a wide range of activation thresholds, consistent with scale-invariant, near-critical dynamics ^26,27^. Ultimately, these results show that structural strength is a primary determinant of where large-scale neuronal avalanches begin and end.

## Author contribution

M.D. M.D.L. conceptualized the study; M.D.L., E.T.L., E.G. and G.C. performed experiments; M.D., G. D., M.A. and M.D.L., analyzed the data; G.S., M.Q., C.G., and P.S. supervised the study; M.D.L., M.D. wrote the manuscript; G.S., M.A., P.S. and D.D. reviewed and edited the manuscript.

## Data Availability Statement

Data are available on reasonable request to the corresponding author.

## Informed Consent Statement

Informed consent was obtained from all subjects involved in the study.

